# The expression of virulence increases outer-membrane permeability and sensitivity to envelope stress in *Salmonella* Typhimurium

**DOI:** 10.1101/2021.06.08.447568

**Authors:** Malgorzata Sobota, Pilar Natalia Rodilla Ramirez, Alexander Cambré, Tiphaine Haas, Delphine Cornillet, Andrea Rocker, Abram Aertsen, Médéric Diard

**Author notes:** Contributed equally (Malgorzata Sobota, Pilar Natalia Rodilla Ramirez, Alexander Cambré).

## Abstract

Environmental cues modulate the expression of virulence in bacterial pathogens. However, while cues that upregulate virulence are often intuitive and mechanistically well understood, this is less so for cues that downregulate virulence. In this study, we noticed that upregulation of the HilD virulence regulon in *Salmonella* Typhimurium (*S*.Tm) sensitized cells to membrane stress mediated by cholate, Tris/EDTA or heat. Further monitoring of membrane status and stress resistance of *S*.Tm cells in relation to virulence expression, revealed that co-expressed virulence factors embedded in the envelope (including the Type Three Secretion System 1 and the flagella) increased permeability, and stress sensitivity of the membrane. Importantly, pretreating the bacteria by sublethal stress inhibited virulence expression and restored stress resistance. As such, these results demonstrate a trade-off between virulence and stress resistance, which explains the downregulation of virulence expression in response to harsh environments in *S*.Tm.

## Introduction

Bacteria constantly sense their environment and adapt to changes by modulating gene expression accordingly. In entero-pathogenic bacteria, physiological and environmental stimuli drive the expression of virulence genes and the outcome of the interaction with the host ^**1-3**^. Virulence can maximize the fitness of pathogens via host exploitation ^**4,5**^. On the other hand, virulence factors can be costly to produce, therefore requiring fine-tuned regulation to ensure balanced and timely expression ^**6**^. However, as intuitive as it sounds, causal links between environmental stimuli, fitness cost of virulence and adaptive evolution of regulatory pathways remain to be demonstrated. Here we address such link in *Salmonella enterica* Typhimurium (*S*.Tm).

*S*.Tm is a facultative intracellular entero-pathogen able to prosper in the intestinal lumen of a broad range of hosts ^**7**^. At the center of the regulatory network of virulence expression sits the AraC-like transcription factor HilD that controls about 250 genes constituting the HilD regulon ^**8-10**^. The expression of the HilD regulon depends on various environmental parameters ^**2**^. Upon entry into stationary phase in rich medium, HilD mediates the OFF to ON virulence switch in a subset of cells (bimodal expression ^**11,12**^). In these conditions, the HilD regulon comprises the *Salmonella* Pathogenicity Island-1 (SPI-1) - Type Three Secretion System (T3SS-1) and associated effectors, the SPI-4 - T1SS and the giant adhesin SiiE, as well as motility and chemotaxis apparatuses ^**9**^ (**Fig. S1, Table S1**). In the gut, these functions prime *Salmonella* to swim toward, attach to and invade enterocytes ^**7,13,14**^. The resulting intestinal innate immune response favors the growth and the transmission of *S*.Tm ^**5,15**^, demonstrating how virulence improves *S*.Tm fitness at the population level^**16**^

However, the fitness-cost of virulence at the single-cell level and the facultative intra-cellular lifestyle of *S*.Tm strongly constrain the expression of the HilD regulon. The expression of invasion factors controlled by HilD correlates with a two-fold reduction of the growth rate ^**12**^. Although such a substantial fitness-cost could threaten the evolutionary stability of virulence in *S*.Tm ^**17**^, the heterogeneous “all or nothing” (ON/OFF) bimodal expression pattern (**Fig. S1A**, ^**11**^) mitigates the cost by allowing only a fraction of *Salmonella* cells to engage in the costly virulence program ^**17**^. Furthermore, once in the *Salmonella* containing vacuole, the invasion genes are silenced, whereas the expression of a specific set of genes ensures the intra-cellular survival of *S*.Tm ^**18,19**^.

A number of signals modulate virulence expression in *S*.Tm. Availability of carbon sources, amino acids, divalent cations, phosphate and oxygen tilt the balance toward inhibition or activation of the HilD regulon (reviewed in ^**2,3**^). Envelope stress, e.g., exposure to low pH, cationic antimicrobial peptides, bile and heat, as well as misassembled outer-membrane proteins, generally inhibits T3SS-1 expression ^**20-23**^. The PhoP-PhoQ two-component system plays a central role in repressing invasion genes upon exposure to low Mg^2+^, low pH and cationic antimicrobial peptides ^**24-26**^. Some of these stimuli are encountered within host cells, making sensing by PhoPQ essential to optimize the shift from invasion to intra-cellular survival ^**27**^. However, it is not clear why sensing bile and excessive heat ^**22,23**^, or misassembled periplasmic outer-membrane proteins by the Rcs system ^**20**^, should inhibit the expression of invasion genes. In fact, the evolutionary causes of the regulation of invasion factors at the population level outside host cells remain to be determined. As such, we do not fully understand what conditions the fact that some *S*.Tm cells should express invasion factors or not, how virulence gene expression intertwines with the general physiology of the bacteria and to what extent this has shaped regulatory networks in *S*.Tm.

Since most HilD controlled invasion factors (i.e., T3SS-1, SPI-4 T1SS, flagella and chemotactic receptors) are embedded within the envelope, we hypothesized that the expression of the HilD regulon could render *S*.Tm intrinsically more sensitive to envelop stress, and that inhibiting the expression of HilD controlled genes could correspondingly be integral part of a general envelop stress response in *S*.Tm. This hypothesis was addressed by comparing outer membrane permeability and survival rate to envelope stresses in populations of *S*.Tm strains expressing the HilD regulon or not. Our results reveal a trade-off between membrane robustness and virulence expression in *Salmonella* with potential consequences for the design of anti-virulence strategies against this pathogen.

## Results

### Proteomic analysis of the HilD regulon on sorted *S*.Tm cells

In this study, we used the ectopic chromosomal P*prgH*::*gfp* reporter (in which GFP expression is controlled by the promoter of the T3SS-1 *prg* operon ^**11**^) as a proxy for the expression of the HilD regulon in *S*.Tm SL1344 (further referred to as wild-type or WT). The distribution of the GFP expression at early stationary phase in LB was clearly bimodal with ca. 63% of the population in the OFF state and ca. 37% in the ON state (**Fig. S1A**). As previously reported, the ON/OFF ratio results from the expression of HilD, whose activity is controlled by the negative regulator HilE at the post-translational level ^**12,17,28**^. Accordingly, the proportion of ON cells was increased to ca. 50% in the Δ*hilE* mutant. Note that the ON/OFF ratio in WT and its Δ*hilE* derivative varied between experiments but the latter consistently yielded more ON cells than the WT (**Fig 1A, 2D, 3A**). Deletion of *hilD*, on the other hand, clearly locked the cells in the OFF state (**Fig. S1A**).

**Fig. 1:**
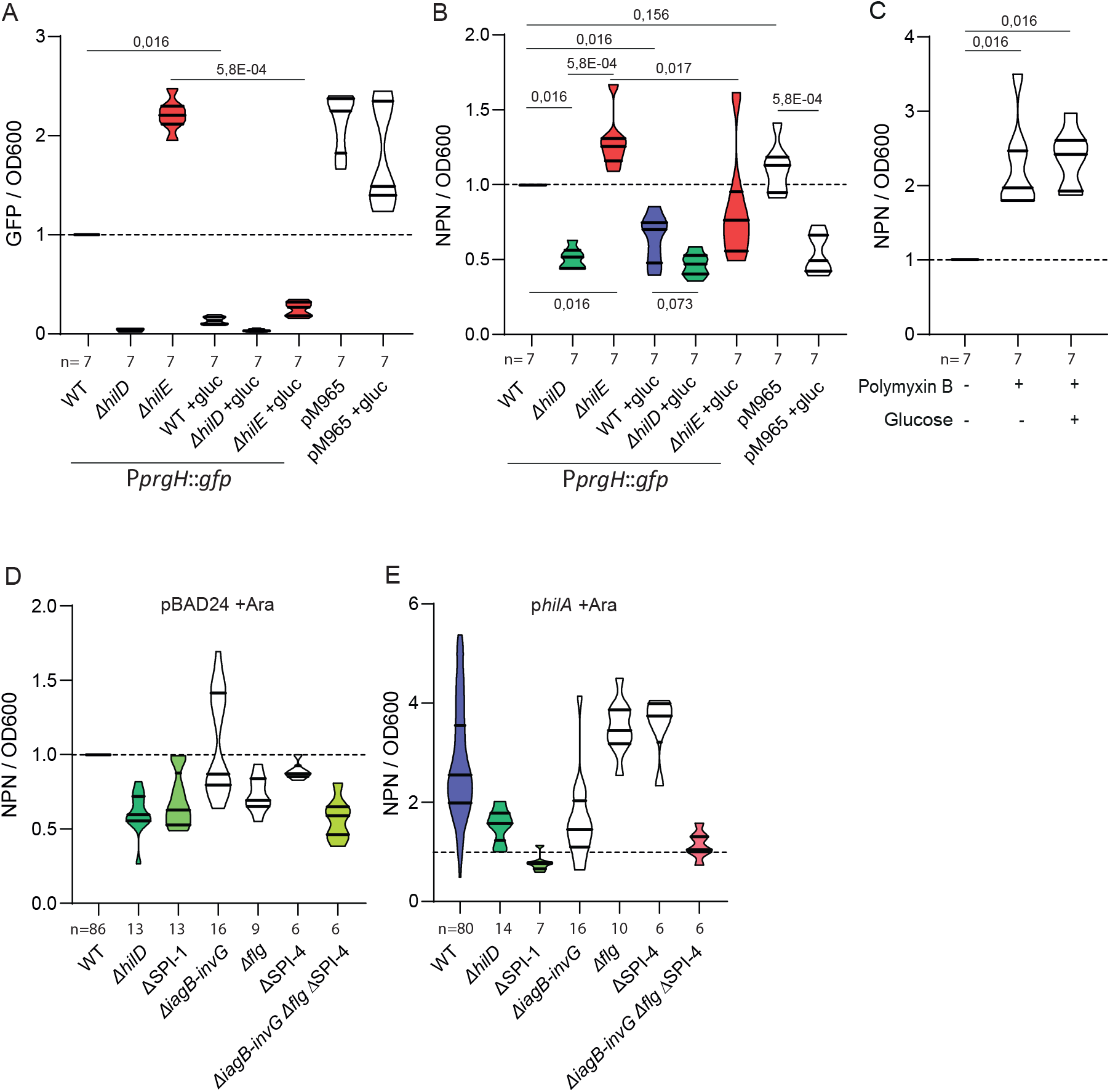
Expression of the HilD regulon increases the permeability of the outer-membrane to NPN. Green fluorescence (GFP from P*prgH*::*gfp*) **(A)** and NPN fluorescence values **(B)** were measured on cells at late exponential phase treated with 10µM NPN, and were divided by the optical density (OD600). Values of each repetition for mutant strains are normalized according to a parallel experiment on the wild type (WT). When indicated, 0.1% of glucose was added to day cultures in order to repress virulence expression (+gluc). A control strain constitutively expressing GFP from pM965 carrying P*rpsM*::*gfp* was used to control for the effect of GFP on the fluorescence readout for NPN uptake. A 2.5µg/mL Polymyxin B treatment to permeabilize the membrane was used as positive control **(C). D,E**. NPN fluorescence values divided by the OD600 measured on day cultures of WT, Δ*hilD*, ΔSPI-1, Δ*iagB-invG*, Δ*flg*, ΔSPI-4 and triple mutant Δ*iagB-inv*G Δ*flg* ΔSPI-4 carrying either pBAD24 **(D)** or p*hilA* **(E)** and treated with 10µM NPN. These values were normalized to the reference WT pBAD24 grew in presence of 1mM arabinose. Values obtained in the absence of arabinose are shown in **figure S2**. For comparisons against the WT, p values were calculated on the raw data using paired Wilcoxon tests. For comparisons between mutants or conditions, p values were calculated using the normalized data and unpaired Mann-Whitney tests. The **table S2** shows p values for comparisons between groups from panels **D** and **E**.

**Fig. 2:**
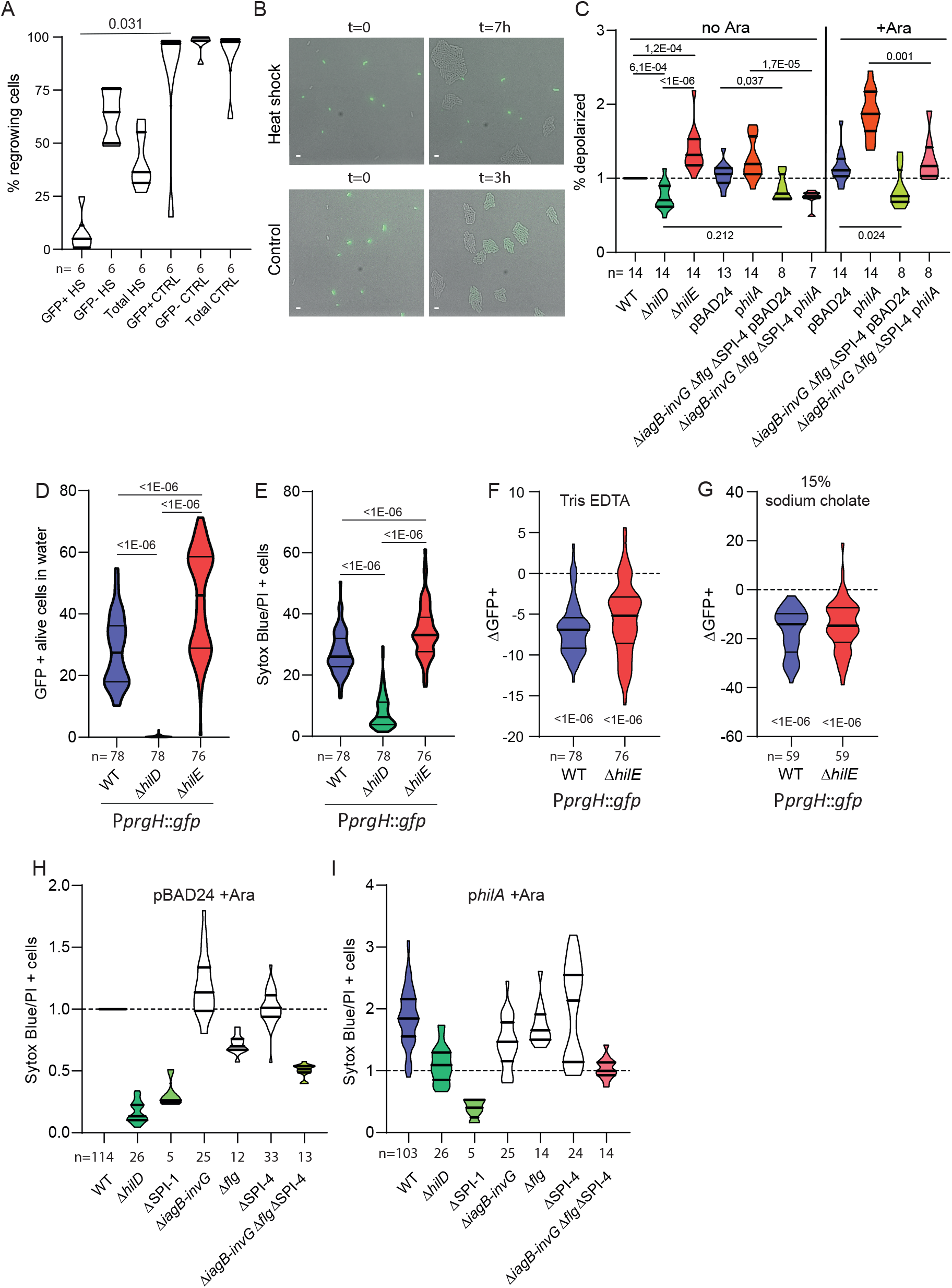
Expression of the HilD regulon comes with membrane stress sensitivity. **A**. Time-lapse microscopy analysis of WT reporter strain (P*prgH::gfp*, GFP) after heat shock (HS, 51°C, 15min) and untreated control (CTRL). Violin plots represent fraction of cells enable to form microcolonies among cells expressing the HilD regulon (GFP+), not expressing the HilD regulon (GFP-) and the total population. **B**. Representative pictures from the time-lapse microscopy experiment. Cells in the upper panel were heat shock treated. Left picture shows cells at t=0, right picture shows cells after 7h. Cells in the lower panel are untreated control. Left picture shows cells in at t=0, right picture shows cells after 3h. Scale bar: 2µm. **C**. Cells at late exponential phase grown in LB supplemented (+ Ara) or not (no Ara) with 1mM L-arabinose to induce the over-expression of *hilA* from p*hilA* (derivative of pBAD24) were stained using 30µm DiOC_2_(3) in the presence of 10mM Tris-1mM EDTA and analyzed by flow cytometry. Unstained WT control and fully depolarized WT cells treated with 100µm CCCP and 30µm DiOC_2_(3) were used to define the depolarized population in each sample (**Fig. S3**). The proportion of depolarized cells was then normalized according to values obtained in parallel experiments using the WT. For comparisons against the WT, p values were calculated on the raw data using paired Wilcoxon tests. For comparisons between mutants or conditions, p values were calculated using the normalized data and unpaired Mann-Whitney tests. **D**. Proportion of GFP expressing cells from day cultures of reporter strains (P*prgH::gfp*, GFP) WT, Δ*hilD* and Δ*hilE*, measured by flow cytometry, unstained by sytox blue or propidium iodide (PI), i.e., alive after exposure to distilled water. **E**. Frequency of cells stained with either sytox blue or PI after treatment with 100mM Tris-10mM EDTA measured by flow cytometry for day cultures of reporter strains WT, Δ*hilD* and Δ*hilE*. **F, G**. Reduction of the GFP positive fraction (ΔGFP+) among WT or Δ*hilE* alive cells treated with 100mM Tris-10mM EDTA compared to distilled water control **(F)** or 15% sodium cholate compared to PBS control **(G)**. Significance of the deviation of the median from 0 estimated by Wilcoxon signed rank test. **H, I**. Normalized frequency of late exponential phase cells stained with either sytox blue or PI after treatment with 100mM Tris-10mM EDTA of Δ*hilD*, ΔSPI1, Δ*iagB-invG*, Δ*flg*, ΔSPI-4, Δ*iagB-invG* Δ*flg* ΔSPI-4 strains harboring pBAD24 **(H)** or p*hilA* **(I)**. Data normalized to parallel WT controls. On the left panel, strains were carrying the empty vector pBAD24, on the right panel p*hilA*. The day cultures were supplemented with 1mM arabinose to induce HilA expression in the strains carrying p*hilA* **(I)**. For comparisons against the WT, p values were calculated on the raw data using paired Wilcoxon tests. For comparisons between mutants or conditions, p values were calculated using the normalized data and unpaired Mann-Whitney tests. The **table S4** shows p values for comparisons of data from panels **H** and **I**.

**Fig. 3:**
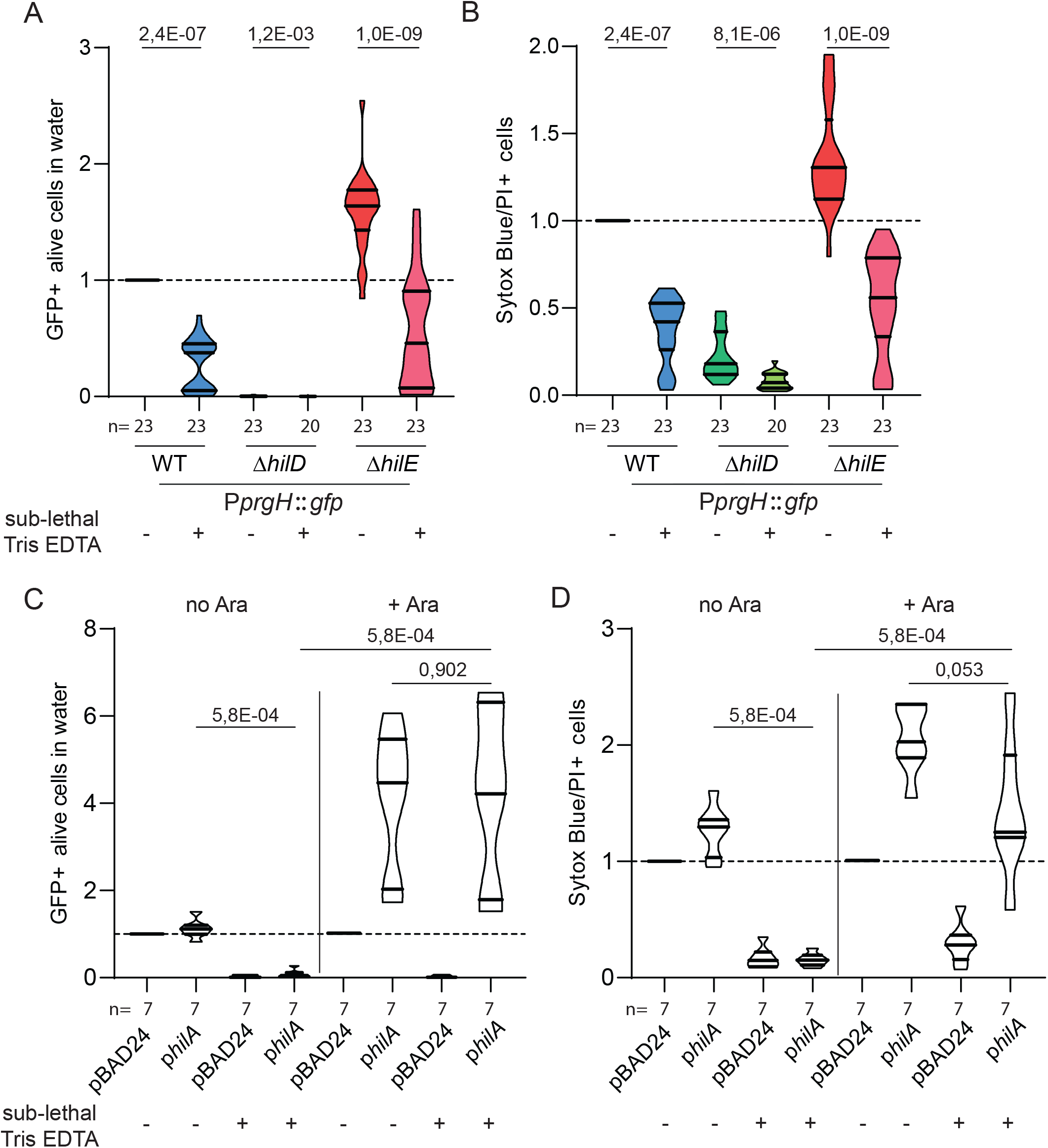
Sub-lethal stress inhibited expression of virulence and increased resistance against lethal Tris EDTA (TE) exposure. Flow cytometry analysis. **A**. Proportion of GFP (P*prgH::gfp*) expressing cells unstained by sytox blue or propidium iodide (PI) (alive) in distilled water of reporter strains WT, Δ*hilD* and Δ*hilE*. Values normalized to WT. When indicated, 4mM Tris-0.4mM EDTA was added to the day cultures as sub-lethal pre-treatment. **B**. Frequency of cells stained with either sytox blue or PI after treatment with 100mM Tris-10mM EDTA in WT, Δ*hilD* and Δ*hilE* strains. Values were normalized to WT. When indicated, 4mM Tris-0.4mM EDTA was added to the day cultures. **C**. Proportion of GFP expressing cells, unstained by sytox blue or PI, in distilled water. WT strain carrying the empty vector pBAD24 or the p*hilA* plasmid allowing for over-expression of HilA. When indicated, 1mM arabinose and/or 4mM Tris 0.4mM EDTA was added in the day cultures. **D**. Frequency of late exponential phase cells stained with either sytox blue or PI after treatment with 100mM Tris-10mM EDTA. When indicated, 1mM arabinose and/or 4mM Tris-0.4mM EDTA were supplemented in the medium. **C and D**. Data normalized by WT pBAD24 (no Ara or +Ara). For comparisons against the WT, p values were calculated using the raw data and paired Wilcoxon tests. For comparisons between mutants or conditions, p values were calculated using data normalized using corresponding WT or WT pBAD24 (no Ara or +Ara) as reference and unpaired Mann-Whitney tests.

To confirm the co-expression of the P*prgH*::*gfp* reporter with HilD regulated genes, we compared the proteomic profile of WT *S*.Tm cells sorted according to the bimodal pattern of the green fluorescence signal (**Fig. S1B, Table S1**). We observed an increase in SPI-1, SPI-4, flagella and chemotaxis protein expression in GFP positive cells, which was consistent with previously published transcriptomic data describing the HilD regulon ^**8-10**^. This validated the use of the P*prgH*::*gfp* fusion as reporter for HilD regulon expression at the single-cell level.

### Expression of HilD increases the permeability of the outer-membrane to a lipophilic compound

Several functions controlled by HilD are large protein complexes embedded in the envelope of *S*.Tm (T3SS-1, SPI-4 T1SS, flagella and chemotactic receptor clusters (**Fig. S1B, Table S1**)) potentially affecting envelope integrity. In order to assess disruption of the outer-membrane we used N-Phenyl-1-Naphthylamine (NPN), a lipophilic dye that is weakly fluorescent in aqueous environments but becomes highly fluorescent in hydrophobic membranes ^**29**^. In growth conditions triggering expression of the HilD regulon (i.e., early stationary phase in LB) (**Fig. 1A**), the WT and the *hilE* mutant accumulated significantly more NPN than the *hilD* mutant (**Fig. 1B**). In contrast, adding glucose to the media drastically reduced the expression of the HilD regulon (**Fig. 1A**) and NPN uptake (**Fig. 1B**). As a control, Polymyxin B, acting as a detergent, increased NPN uptake independently of glucose presence. We tested the effect of GFP expression (used to monitor induction of the HilD regulon) by constitutively expressing it from a plasmid (pM965; transcriptional fusion P*rpsM*::*gfp*) (**Fig. 1A**), which demonstrated that the presence of GFP itself did not affect NPN uptake or its signal (**Fig. 1B**). We then evaluated the contribution of SPI1, SPI4, and flagella in increasing membrane permeability in *S*.Tm (**Fig. 1D and E**). Strains carrying pBAD24 and growing in the presence of arabinose (**Fig. 1D**) were compared with strains over-expressing HilA from the pBAD24 derivative p*hilA* (**Fig. 1E**). Of note, results obtained with strains carrying pBAD24 and p*hilA* but growing in the absence of arabinose showed similar patterns of NPN uptake as strains carrying pBAD24 in presence of arabinose (no HilA over-expression, **Fig. S2**). The full SPI-1 deletion (including the *hilD* gene) phenocopied the Δ*hilD* mutant, validating our first observation. The deletion of the *iagB*-*invG* locus in SPI-1 (i.e. removing operons *iag, spt, sic, iac, sip, sic, spa*, and *inv*, but keeping transcriptional regulators *hilD, hilC, hilA*, and *invR* intact), the *flgBCDEFGHIJ* operon (thereafter shortened *flg*), or SPI-4 did not fully restore membrane integrity suggesting that increased NPN permeability in the HilD regulon expressing cells was multifactorial. Indeed, only combining deletions of *iagB*-*invG, flg* and SPI-4 phenocopied the *hilD* mutant (**Fig. 1D**).

Increasing T3SS-1 and SPI-4 production via overexpression of the transcriptional factor HilA further confirmed these results (**Fig. 1E**). Arabinose induced overexpression of HilA led to drastic increase in NPN uptake in the WT. In these conditions, membrane permeability was restored by the *iagB*-*invG* deletion and in the triple mutant *iagB*-*invG flg* SPI-4, but unchanged by deleting the *flg* operon (not under the control of HilA) or SPI-4. The reproducible pattern of NPN uptake in various mutants when HilA is not over-expressed (**Fig. 1D and S2**), suggests that the flagella was the most important contributor followed by the T3SS-1, while the SPI-4 T1SS did not play a significant role in increasing outer-membrane permeability to NPN. The **table S2** gathers statistical analysis results from this dataset.

### Resistance to outer-membrane disrupting treatments is reduced in HilD expressing cells

Given the higher permeability of the outer-membrane to the lipophilic compound NPN in *hilD* expressing populations, we hypothesized that the expression of the HilD regulon could be associated with lesser resistance to stress further disturbing the outer-membrane.

Heat provokes outer-membrane permeability ^**30**^. Therefore, we first measured survival of *S*.Tm exposed to a mild heat shock (HS, 51°C, 15 min) at the single-cell level with time-lapse fluorescence microscopy. The cells were observed during 16h post-treatment. This revealed that a vast majority of cells expressing the HilD regulon (i.e. GFP positive) was unable to resume growth (**Fig. 2A and B** heat shock upper panels), while the rest of the population regrew after a lag period. Untreated cells were able to grow normally regardless of their HilD expression state, with ON cells shifting OFF and diluting the GFP (**Fig. 2B** control lower panels).

We then measured membrane polarization in cells exposed to 10mM Tris-1mM EDTA, which increases Gram negative bacteria permeability to the dye 3,3′-diethyloxa-carbocyanine iodide (DiOC_2_(3)) (**Fig. 2C**) ^**31**^ and destabilize the outer-membrane ^**32**^. Permeable but polarized cells tend to accumulate DiOC_2_(3) to a point of dye aggregation shifting its fluorescence from green (all stained cells) to red (polarized cells). We followed the gating strategy described in **figure S3** to estimate the proportion of depolarized cells (green) among the population of stained cells (red and/or green) in WT and mutant strains. Exposure to the proton ionophore carbonyl cyanide 3-chlorophenylhydrazone (CCCP) was used as control condition in which the membrane potential is abolished and most DiOC_2_(3) stained cells remained green (depolarized). Unstained cells and cells exposed to CCCP and DiOC_2_(3) served as references (**Fig. S3**).

A pattern consistent with NPN uptake emerged from these experiments. The WT strain and the *hilE* mutant showed significantly higher proportion of depolarized cells compared to the *hilD* mutant. The phenotype was enhanced by HilA overexpression from p*hilA* in the presence of arabinose and rescued in the *iagB*-*invG flg* SPI-4 triple mutant (**Fig. 2C**). In the context of endogenous HilA expression, the phenotype of the triple mutant TTSS-1/flagella/SPI-4 phenocopied the *hilD* mutant.

Membrane polarization in absence of stress was measured in control experiments using 3,3’-Dipropylthiadicarbocyanine Iodide (DiSC_3_(5)) (**Fig. S4**). This red fluorescent hydrophobic probe accumulates in the membrane of polarized cells ^**33**^. This assay was compatible with the readout for expression of the HilD regulon with the reporter P*prgH*::*gfp*. We observed no difference in DiSC_3_(5) staining in HilD ON (GFP+) vs OFF cells (GFP-) (**Fig. S4**) suggesting that HilD expressing cells were not inherently depolarized or hyperpolarized. The membrane depolarization observed with DiOC_2_(3) was therefore due to exposure to 10mM Tris-1mM EDTA further destabilizing the outer-membrane, especially when *S*.Tm co-express the TTSS-1 and the flagella (**Fig. 2C**).

Thirdly, we measured sensitivity to a 100mM Tris-10mM EDTA (TE) lethal treatment that provokes lysis of Gram-negative bacteria by destabilizing the lipopolysaccharide ^**32**^. We used two complementary approaches, flow cytometry and microscopy, to quantify the proportion of dead cells detectable after TE stress and the fraction of ON (GFP+) cells among the survivors. Flow cytometry can measure fluorescence in a relatively high number of cells and allows testing multiple conditions in a single run. We used GFP as readout for HilD expression and two dyes to stain the dead cells: propidium Iodide (PI) or sytox blue. Thoroughly filtered media and buffers ensured that no debris could interfere with the quantification. **Figure 2D** shows the expression of the HilD regulon expression (proportion of GFP+ cells exposed to distilled water, used as solvent generating hypo-osmotic shock). After TE treatment, we observed a clear increase in the proportion of dead cells in wild-type and the *hilE* mutant compared to the *hilD* mutant with both dyes (**Fig. 2E**). Although the Sytox Blue had the tendency to stain slightly more cells than the PI (**Fig. S5A**), we judged the overlap sufficient to pool both staining results, obtained in parallel technical replicates, in every live-dead datasets.

In addition, the proportion of ON cells decreased in the surviving populations compared to the water only control (**Fig. 2F**). The timing of the experiment (30’ of treatment before cytometry analysis) was too short to allow ON cells to switch OFF and to dilute the GFP by dividing. Expressing GFP from a constitutive promoter did not change the fraction of dead cells detected by Sytox blue or PI staining (**Fig. S5B**). We ruled out the contribution of non-fluorescent and unstained debris formed during stress by quantifying the fraction of non-fluorescent events from stressed cells constitutively expressing GFP (**Fig. S5C**). Moreover, a treatment with 15% sodium cholate, a natural detergent present in bile, was even more potent at reducing the proportion of ON cells in wild-type (**-14%** vs **-6.8%** with TE) and the *hilE* mutant (**-14.7%** vs **-5.2%** with TE) (**Fig. 2G**).

Observing WT reporter bacteria under the microscope after TE treatment confirmed that, although ca. 70% of visible cells were able to resume growth whatever their status (HilD regulon ON (GFP+) or OFF (GFP-)) (**Fig. S6A**), the proportion of ON cells was significantly reduced (17% less ON cells compared to control (**Fig. S6B**)). This indicates that TE provoked lysis of a substantial amount of bacteria, with a higher probability for the ON cells to lyse. Counting colony-forming units (CFU) post-treatment confirmed that Sytox blue or PI staining and counting the re-growing cells under the microscope underestimated the fraction of cells dying upon TE treatment (**Table S3**). Nevertheless, flow cytometry and microcopy showed comparable results with 30% of WT cells stained by Sytox blue/PI (**Fig. 2E**) or not re-growing on the agar pad after TE treatment (**Fig. S6A**).

Cumulative *iagB*-*invG flg* SPI-4 deletions reduced sensitivity to TE treatment, although they did not consistently phenocopy the resistance of the *hilD* or SPI-1 deletion mutants (comprising *hilD*) (**Fig. 2H and I**). As observed for NPN uptake, the deletion of SPI-4 did not markedly affect sensitivity to TE. However, the individual deletion of the *flg* operon significantly reduced TE sensitivity (**Fig 2H**). The over-expression of HilA (p*hilA* + Arabinose) drastically enhanced TE sensitivity (**Fig. 2I**). The full SPI-1 deletion and, to a lesser extent the deletion of *hilD* alone, of Δ*iagB-invG*, and of *iagB*-*invG flg* SPI-4 restored resistance. In this case again, *flg* and SPI-4 deletions had no effect because not induced. Controls in the absence of arabinose reflects the pattern observed for strains carrying pBAD24 in the presence of arabinose (**Fig. S7**). **Table S4** gathers statistical analysis results from this dataset.

### Sub-lethal stress inhibits expression of virulence and increases resistance to lethal exposure to Tris-EDTA (TE)

Given that the expression of the HilD regulon sensitizes *S*.Tm to heat, sodium cholate and TE exposure, and previous observations that bile or heat shock inhibit T3SS-1 expression ^**20-23**^, we reasoned that reacting to sub-lethal stress exposure by shutting down the HilD regulon should rescue *S*.Tm against lethal stress. To address the effect of stress on HilD regulon expression and survival to subsequent lethal exposure, we grew the bacteria in LB medium containing TE at sub-lethal concentration (4mM Tris-0.4mM EDTA) before exposure to lethal dose (100mM Tris-10mM EDTA). As expected, the proportion of ON cells was strikingly reduced when *S*.Tm grew in the presence of TE at sub-lethal concentration (**Fig. 3A**) and survival was increased when the bacteria were then exposed to lethal dose (**Fig. 3B**). Overexpression of *hilA* restored P*prgH*::*gfp* expression (**Fig. 3C**) and with it, the high death rate in sub-lethal TE pretreated cells exposed to lethal TE concentration (**Fig. 3D**). This suggested that the expression of genes downstream of *hilA* increased stress sensitivity even if stress response is triggered by sub-lethal stress exposure. This demonstrated that shutting down the expression of the HilD regulon under stressful conditions increased chances of surviving harsher environments otherwise lethal for the ON cells.

## Discussion

In *S*.Tm, the cost of virulence expression drives within-host emergence of attenuated mutants harboring loss-of-function mutations in positive transcriptional regulators of virulence (e.g. *hilD* and *hilC* ^**17,34**^). Such attenuated mutants consistently emerge during chronic infections in mice and loss-of-function mutations in *hilD* were detected in a large collection of sequenced natural isolates at a frequency suggesting positive selection ^**35**^. The genetic instability of virulence in *S*.Tm could be exploited to fight against this pathogen increasingly resistant to antibiotics ^**36**^. However, the tight regulation of bimodal expression of virulence impairs fixation of attenuated mutants during within-host growth, ensuring transmission of the virulent genotype ^**17**^. Understanding the cost of virulence, and how it relates to expression regulation, could allow identifying and modulating ecological factors in order to drive the evolution of *S*.Tm toward attenuation.

Until now, the only identified cost of virulence in *S*.Tm was a two-fold reduction of the growth rate in cells expressing the HilD regulon ^**12**^. The molecular mechanism underlying this growth defect is still unclear. Here, we discovered that the expression of HilD increased membrane permeability to the hydrophobic compound NPN (**Fig. 1**) and sensitized *S*.Tm to stresses that disrupt the envelope (i.e., short exposure to mild heat, Tris/EDTA and sodium cholate, **Fig. 2**). To identify the functions involved, we compared mutants lacking different operons upregulated by HilD coding for the T3SS-1, the flagella and the SPI-4 T1SS. The deletion of T3SS-1 components partially restored stress resistance. This might relate to the fact that mislocalized T3SS’s components, like the secretin, can disrupt the membrane, as reported in *Yesinia enterocolitica* and *E. coli* mutants that lack functional phage-shock proteins ^**37,38**^, as well as in *Pseudomonas aeruginosa* ^**39**^. However, we found that the flagella also played a role, in accordance with the observation that flagella expression increases intrinsic death rate in *E. coli* MG1655, a strain that does not possess a T3SS ^**40**^. Virulence-associated envelope fragility in *S*.Tm was therefore clearly multifactorial although the SPI-4 T1SS was not significantly involved.

The relative contribution of envelope destabilization by multi protein complexes (T3SS-1 and flagella) and of the energetic burden of secretion and motility to the fitness-cost of virulence is difficult to assess as these processes inevitably intertwine. Nevertheless, the comparative proteomic analysis of untreated HilD ON *vs*. OFF cells did not reveal any significant extra-cytoplasmic stress response in the ON cells, such as the induction of the RpoE or Cpx regulons ^**41**^. Therefore, the bacteria expressing the HilD regulon did not seem more intrinsically stressed than the OFF cells. As previously shown in *Pseudomonas putida* expressing the flagella ^**42**^, an energetic cost of virulence can impair the ability to cope with certain stresses. However, membrane polarization was comparable between HilD ON and OFF *S*.Tm cells.

The repression of the HilD regulon by sub-lethal exposure to Tris-EDTA was in accordance with known inhibition of T3SS-1 expression in *S*.Tm exposed to envelop stress (e.g. heat, bile or CAMP ^**21-23**^), and deprivation of divalent cations sensed by the two-component system PhoPQ ^**24**^. The artificial over-expression of HilA showed that decoupling stress response from inhibition of virulence expression was detrimental to *S*.Tm in harsh environment. To the best of our knowledge, this is the first dataset demonstrating a link between regulation of virulence expression and stress response in *S*.Tm driven by the fitness-cost of virulence. We propose that such trade-off between virulence and membrane robustness in *S*.Tm shaped the regulation of virulence expression by integrating it into the general stress response. Moreover, since many virulence determinants are typically deployed at the level of the cell envelope, this trade-off and regulatory integration might constitute a more general paradigm present as well in other bacterial pathogens.

To conclude, we demonstrate that the cost of virulence in *S*.Tm is pleiotropic but strongly envelope-related, and that such virulence costs can be mitigated by stress response regulation. To allow for precise measurements and reasonable throughput, our results were obtained in controlled *in vitro* settings. Nevertheless, although beneficial to *S*.Tm ^**5,15**^, the intestinal inflammation generates a harsh environment ^**43**^ potentially more lethal to cells expressing the virulence program controlled by HilD. Further work should reveal if impaired membrane robustness constitutes in itself a significant selective force that could promote the rise of avirulent mutants within infected hosts ^**17,34**^. Alternatively, biological stress acting more specifically on the HilD expressing cells could be identified and harnessed for the design of novel anti-virulence strategies.

## Materials and methods

### Bacterial cultivation

All the strains and plasmids used in this study are listed in **table S5**. *Salmonella enterica* serovar Typhimurium SB300 (SL1344)^**44**^ and derivatives was cultivated at 37°C using LB liquid or solid media (Difco). Antibiotic selection was performed with 100µg/ml ampicillin (Sigma), 6 µg/ml chloramphenicol (Sigma), 50µg/ml kanamycin (Sigma) when needed. The day cultures were prepared by 1:100 dilution of overnight cultures in 2mL LB and incubated for 4h at 37°C with shaking. When stated, media were supplemented with 0.1% w/v D-glucose (Sigma) to inhibit the expression of the HilD regulon. For induction of genes under the inducible P*ara* promoter (in p*hilA*), cultures were supplemented with 1mM L-arabinose. Sub-lethal pre-treatment of cells was performed using a mixture of 4mM Tris (Sigma) and 0.4mM EDTA (Sigma) supplemented in the day culture.

### Mutant constructions

*Salmonella* mutants used in this study were constructed by homologous recombination using the λ-Red gene replacement system as described in ^**45**^. To enable selection of recombinants the chloramphenicol acetyltrasferase (*cat*) gene was amplified from the pKD3 plasmid template using primers containing 40bp region homologous to flanking regions of the target gene in the *Salmonella* chromosome. The PCR product was transformed by electroporation into the recipient strain harboring pKD46 helper plasmid encoding λ phage *red, gam* and *exo* genes under an arabinose inducible promoter. Recombinant bacteria were selected on LB plates containing chloramphenicol. Following λ-Red recombination, the chloramphenicol resistance cassette was cured using the flipase encoded on the pCP20 helper plasmid. Correct gene replacement and resistance cassette deletions were confirmed by PCR. All primers used in this study are listed in **table S6**. Bacteriophage P22 HT/int mediated transduction was used to transfer the constructions in the desired background ^**46**^.

### Stress resistance analysis by flow cytometry

10µl of 4h day culture in LB was added into 90µl of stress media or vehicle combined with dead staining. Stress media contained either 15% (w/v) sodium cholate in Phosphate Buffer Saline (PBS) or 100mM Tris-10mM EDTA in ddH^2^O. Dead staining was either propidium iodide (PI) (Invitrogen) at a final concentration of 30µg/ml or sytox blue (Invitrogen) at a final concentration of 10µM. Stress exposure was performed in flat bottom 96 well plates incubated for 30min at 37°C. After incubation, cells were diluted 10X in filtered PBS and analyzed by flow cytometry using LSR Fortessa (BD Bioscience) operated with the FACS Diva software (BD Bioscience). Data acquisition was performed until 50.000 events of unstained cells were recorded using excitation with 561nm laser and band pass filter 610/20nm for PI, and excitation with 405nm laser and band pass filter 450/50nm for Sytox Blue. The GFP signal was recorded using excitation with 488nm laser and band pass filter 512/25nm and 505LP. Data were processed using FlowJo V10 software (FlowJo, LCC). Events were first gated for unstained cells (PI/sytox blue negative), and further selected for GFP positive cells when the reporter strain was used.

### Membrane potential analysis by flow cytometry

1ml of 4h day cultures were centrifuged for 3min at 9500rpm in a tabletop centrifuge. Supernatants were discarded and cell pellets were re-suspended in 1ml of 10mM Tris-1mM EDTA before staining by DiOC_2_(3) or PBS before staining by DiSC_3_(5). CCCP treatment and staining were carried out in 96 well plates in a total staining volume of 50µL. 5µL of washed cells were added into 10mM Tris-1mM EDTA or PBS. In the control wells, CCCP was added at a final concentration of 100µM in DMSO (Sigma). Cells were incubated for 30min at 37°C. After incubation, DiOC_2_(3) (Invitrogen) was added to a final concentration of 30µM or DiSC_3_(5) (Invitrogen) was added to the final concentration of 2µM. Both dyes were prepared in DMSO. The cells were incubated for 5min at room temperature. 20µL were transferred into 180µL of 10mM Tris-1mM EDTA (for DiOC_2_(3) staining) or PBS (for DiSC_3_(5) staining) and analyzed by flow cytometry using LSR Fortessa (BD Bioscience) operated by FACS Diva software (BD Bioscience). 50.000 events were recorded using excitation with 488nm laser and band pass 542/27 for green fluorescence and excitation with 488nm laser and band pass 685/35 for red fluorescence with DiOC_2_(3) staining or 640nm laser and band pass 670/14 for red fluorescence with DiSC_3_(5). Data were processed using FlowJo V10 software (FlowJo, LCC).

### Proteomics analysis

#### Culture and sorting

Wild-type SL1344 cells harboring the GFP reporter for HilD regulon expression (fusion P*prgH*::*gfp*) in late exponential phase (grown for 4h at 37°C after 1:100 dilution of an overnight culture in LB) were collected by centrifugation, resuspended and diluted in PBS. Samples were sorted according to green fluorescence intensity using a FACSAria III (BD Biosciences) with scatter and fluorescence channels for GFP, excitation 488nm, with 514/30nm band pass with the precision on yield. The flow cytometer was calibrated with CST beads. During the sorting process, the sample and the collection tubes were kept at 4°C or on ice.

After sorting, cells were collected by centrifugation at 12000rpm for 10min at 4°C. After each centrifugation step, cells resuspended in the remaining supernatant were transferred to smaller tubes: first 5ml and then 1.5ml protein low binding microcentrifuge tubes (Eppendorf). After the final centrifugation step, pellets were stored at -80°C until further processing.

#### Sample Preparation

Frozen sorted cell pellets were lysed in 20µL of lysis buffer (1% Sodium deoxycholate (SDC), 10mM TCEP, 100mM Tris, pH=8.5) using twenty cycles of sonication (30s on, 30s off per cycle) on a Bioruptor (Dianode). Following sonication, proteins in the bacterial lysate were heated at 95°C for 10 min. Proteins were then alkylated using 15mM chloroacetamide at 37°C for 30min and further digested using sequencing-grade modified trypsin (1/25, w/w, trypsin/protein; Promega, USA) at 37°C overnight. After digestion, the samples were acidified with TFA to a final concentration of 1%. Peptides desalting was performed using iST cartridges (PreOmics, Germany) following the manufactures instructions. After drying the samples under vacuum, peptides were stored at -20°C and dissolved in 0.1% aqueous formic acid solution at a concentration of 0.25mg/ml upon use.

#### Mass Spectrometry Analysis

Peptides were subjected to LC-MS analysis using an Orbitrap Fusion Lumos Mass Spectrometer equipped with a nanoelectrospray ion source (both Thermo Fisher Scientific). Peptide separation was carried out using an EASY nLC-1200 system (Thermo Fisher Scientific) equipped with a RP-HPLC column (75μm × 36cm) packed in-house with C18 resin (ReproSil-Pur C18-AQ, 1.9μm resin; Dr. Maisch GmbH, Germany) and a custom-made column heater (60°C). Peptides were separated using a step-wise linear gradient from 95% solvent A (0.1% formic acid, 99.9% water) and 5% solvent B (80% acetonitrile, 0.1% formic acid, 19.9% water) to 35% solvent B over 45 min, to 50% solvent B over 15 min, to 95% solvent B over 2min, and 95% solvent B over 18min at a flow rate of 0.2 µl/min.

The mass spectrometer was operated in DDA mode with a cycle time of 3 seconds between master scans. Each master scan was acquired in the Orbitrap at a resolution of 240,000 FWHM (at 200 m/z) and a scan range from 375 to 1600 m/z followed by MS2 scans of the most intense precursors in the linear ion trap at “Rapid” scan rate with isolation width of the quadrupole set to 1.4 m/z. Maximum ion injection time was set to 50 ms (MS1) and 35 ms (MS2) with an AGC target set to 250% and “standard”, respectively. Only peptides with charge state 2–5 were included in the analysis. Monoisotopic precursor selection (MIPS) was set to Peptide, and the Intensity Threshold was set to 5e3. Peptides were fragmented by HCD (Higher-energy collisional dissociation) with collision energy set to 35%, and one microscan was acquired for each spectrum. The dynamic exclusion duration was set to 30s.

#### Protein Identification and Label-free Quantification

The acquired raw-files were imported into the Progenesis QI software (v2.0, Nonlinear Dynamics Limited), which was used to extract peptide precursor ion intensities across all samples applying the default parameters. The generated mgf-files were searched using MASCOT against a decoy database containing normal and reverse sequences of the *Salmonella typhimurium* (strain SL1344) (UniProt, release date 07.01.2019, in total 10098 sequences) and commonly observed contaminants generated using the SequenceReverser tool from the MaxQuant software (Version 1.0.13.13). The following search criteria were used: full tryptic specificity was required (cleavage after lysine or arginine residues, unless followed by proline); 3 missed cleavages were allowed; carbamidomethylation (C) was set as fixed modification; oxidation (M) and protein N-terminal acetylation were applied as variable modifications; mass tolerance of 10 ppm (precursor) and 0.6 Da (fragments). The database search results were filtered using the ion score to set the false discovery rate (FDR) to 1% on the peptide and protein level, respectively, based on the number of reverse protein sequence hits in the datasets.

Quantitative analysis results from label-free quantification were normalized and statically analyzed using the SafeQuant R package v.2.3.4 (https://github.com/eahrne/SafeQuant/ (PMID: 27345528)) to obtain protein relative abundances. This analysis included global data normalization by equalizing the total peak/reporter areas across all LC-MS runs, summation of peak areas per protein and LC-MS/MS run, followed by calculation of protein abundance ratios. Only isoform specific peptide ion signals were considered for quantification. The summarized protein expression values were used for statistical testing of between condition differentially abundant proteins. Here, empirical Bayes moderated t-Tests were applied, as implemented in the R/Bioconductor limma package (http://bioconductor.org/packages/release/bioc/html/limma.html). The resulting per protein and condition comparison p-values were adjusted for multiple testing using the Benjamini-Hochberg method.

### Time-Lapse Microscopy

For heat shock, 1mL of the overnight culture was centrifuged and cells resuspended in the same volume of 0.85% KCl (Sigma). 70µl of the suspension was transferred into a 250µl PCR tube and incubated in a thermocycler (Biometra) 15min at 51°C. After this, cells were diluted 1/50 in PBS.

For Tris-EDTA treatement (TE), cells in late-exponential phase were obtained by diluting the overnight culture (1:100) in LB and incubation for 4h at 37°C with continuous shaking at 200rpm. Then 100µl of these cells were added to microcentrifuge tubes with 900µl 100mM Tris-10mM EDTA in ddH_2_O (treatment) or ddH_2_O (control) pre-warmed at 37 °C (i.e., 1:10 dilution), and incubated at 37°C for 30 min in a without shaking. After this, the cells were diluted 1/5 in PBS.

Treated and control cells were inoculated onto a thin matrix of LB-agarose attached to a microscope slide. These slides were prepared as follows: a sticky frame (Gene Frame AB-0578) was attached to a standard microscope slide. The resulting cavity was filled with heated LB supplemented with 2% agarose (A4718; Sigma-Aldrich) and covered with a standard microscope slide. After cooling and removal of the cover slide, strips of agarose were removed with the use of a surgical scalpel blade, resulting in two rectangles of LB-agarose (approximately 5×7mm side), one in the center of each half of the frame. Cells (2µl) were spotted in the center of the pads, and the frame was sealed with a coverslip (24×60mm). Each slide contained one control and one treated sample.

Microscope slides were examined using an inverted microscope (DeltaVision Core, Cytiva Life Sciences, mounted on a motorized Olympus IX71 stand) equipped with an environmental chamber (World Precision Instruments) set at 37°C for up to 16h. Images were acquired using a 60x 1.42 NA Plan Apo N objective (Olympus), with the matching 1.524 refractive index immersion oil (Cargille-Sacher Laboratories) and a solid-state illumination (Spectra X light engine, Lumencore).

Images were recorded with a CoolSNAP HQ2 (Photometrics) at each field every 10 minutes using brightfield (filter “POL”, transmittance 32% and exposure time 0.015 s) and fluorescence once per hour to prevent phototoxicity (single band pass emission filter 525/48 nm, transmittance 32% and exposure time 0.025s). Up to 12 fields were selected per condition and acquired using the microscope control software (DeltaVision SoftWoRx Suite 7.1.0, Cytiva Life Sciences). Images have been analyzed by manual counting using Fiji (ImageJ 1.53c). GFP+ and GFP-cells were identified according to a unique threshold based on fluorescence intensity that was applied on the first picture of every series. On average, 565+/-224, 636+/-248, 493+/-117 and 436+/-118 cells were analyzed per experiment in HS, HS control, TE and TE control conditions respectively.

### 1-N-phenylnaphtylamine (NPN) uptake assay

1ml of day culture was centrifuged 3min at 9500rpm in a tabletop centrifuge and the pellet was washed twice with 1mL of PBS.

NPN uptake was measured in a flat bottom 96 well plates (black walls, Costar) with a final NPN (Sigma) concentration of 10μM. 2.5μg/mL Polymyxin B (Sigma) was used as permeabilizing agent in the positive control. In order to achieve an OD ranging from 0.4 to 0.6, 120 μl of the cell suspension was added to each well, then adding up to 200 μl with PBS (with polymyxin B for the positive controls). NPN was added last, and the measurements were performed immediately.

The readout was recorded in a plate reader Synergy H4 Hybrid Reader (BioTek Instruments GmbH) controlled with the Gen5 software. The following excitation/emission wavelengths (in nm) used were 350/420 for the NPN signal and 491/512 for GFP signal. Bandwidth was 9nm in all four cases, the emission was recorded from the top (Optics: Top) with a gain of 100 for GFP and 80 for NPN. OD was measured using 600 nm wavelength. The measurement was performed at 37°C with fast continuous shaking and NPN signal measurements were collected with 1min intervals for 1h. GFP and OD measurements were taken at the beginning and at the end of the run. We considered the fluorescence values after stabilization of the signal. Thus, one time point was selected per experiment, approximately 45 minutes after the start of the measurements (ranging from 35 to 50 minutes).

### Statistical analysis

Statistical analysis was conducted using GraphPad Prism 8.0.2. After normalization to the WT, outliers were identified using the ROUT method with Q=1%. Statistical significance was assessed using the raw data devoid of the outliers via Wilcoxon matched-pairs signed rank test when analyzing comparisons against the WT, and the normalized data devoid of outliers via Mann-Whitney test for comparisons between mutants. *P* values□of <0.05 were considered statistically significant. Violin plots show the empirical distribution of the data, the median and quartiles. The number of independent biological replicates per condition is indicated below the figures (n=x).

## Author contributions

MS, PNRR, AA and MD designed the experiments. MS and PNRR performed NPN uptake experiments for figure 2. MS performed the flow cytometry experiments for figures 1, 2 and 3. PNRR performed time-lapse microscopy for figure 2 and proteomics (fig S1). AC, AR, TH and DC assisted in design and execution of the experiments. MD led the project and wrote the manuscript. All of the co-authors have read and commented on the manuscript.

## Acknowledgements

We would like to acknowledge Stefan Bassler, the FACS core facility, the proteomics core facility and the imaging core facility of the Biozentrum for technical support as well as all the members of Diard laboratory for scientific input on this project.

MD was supported by a SNF professorship (PP00PP_176954). The Biozentrum of the University of Basel provided MS’s PhD stipend.

The group of AA was supported by a fellowship (1135116N to AC) and a grant (G0C7118N) from the Research Foundation Flanders (FWO-Vlaanderen).

## Figure captions

**Fig. S1:**
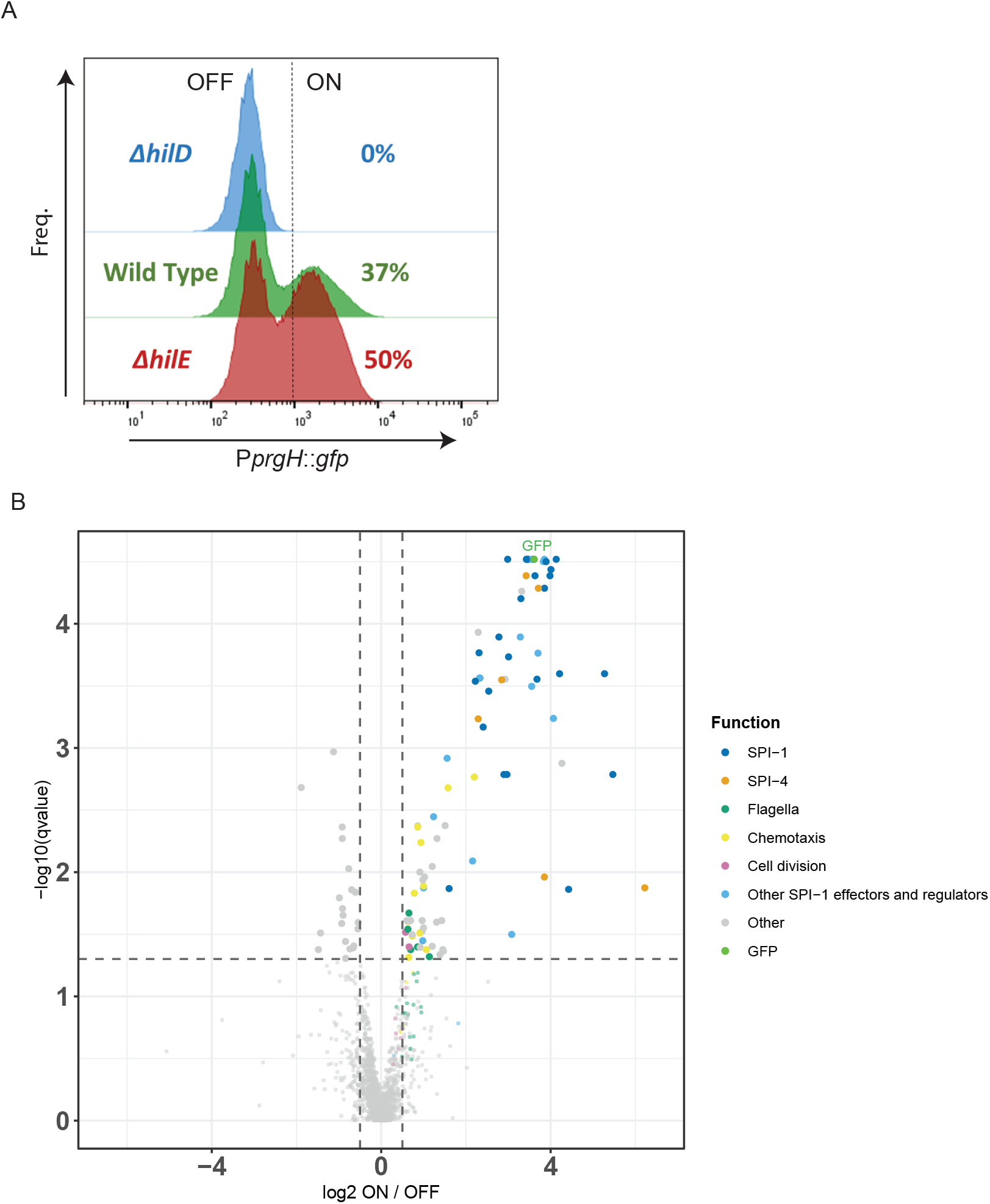
Comparative proteome on *S*.Tm cells sorted according to P*prgH::gfp* expression. **A**. Histogram showing the P*prgH::gfp* reporter expression pattern in strains WT, Δ*hilD* and Δ*hilE*. **B**. Volcano plot showing differential abundance of proteins in sorted GFP+ (ON) and GFP-(OFF) cells of *S*.Tm WT reporter strain. **Table S1** lists these proteins, their functions and provides details about fold-change, statistics and loci.

**Fig. S2:**
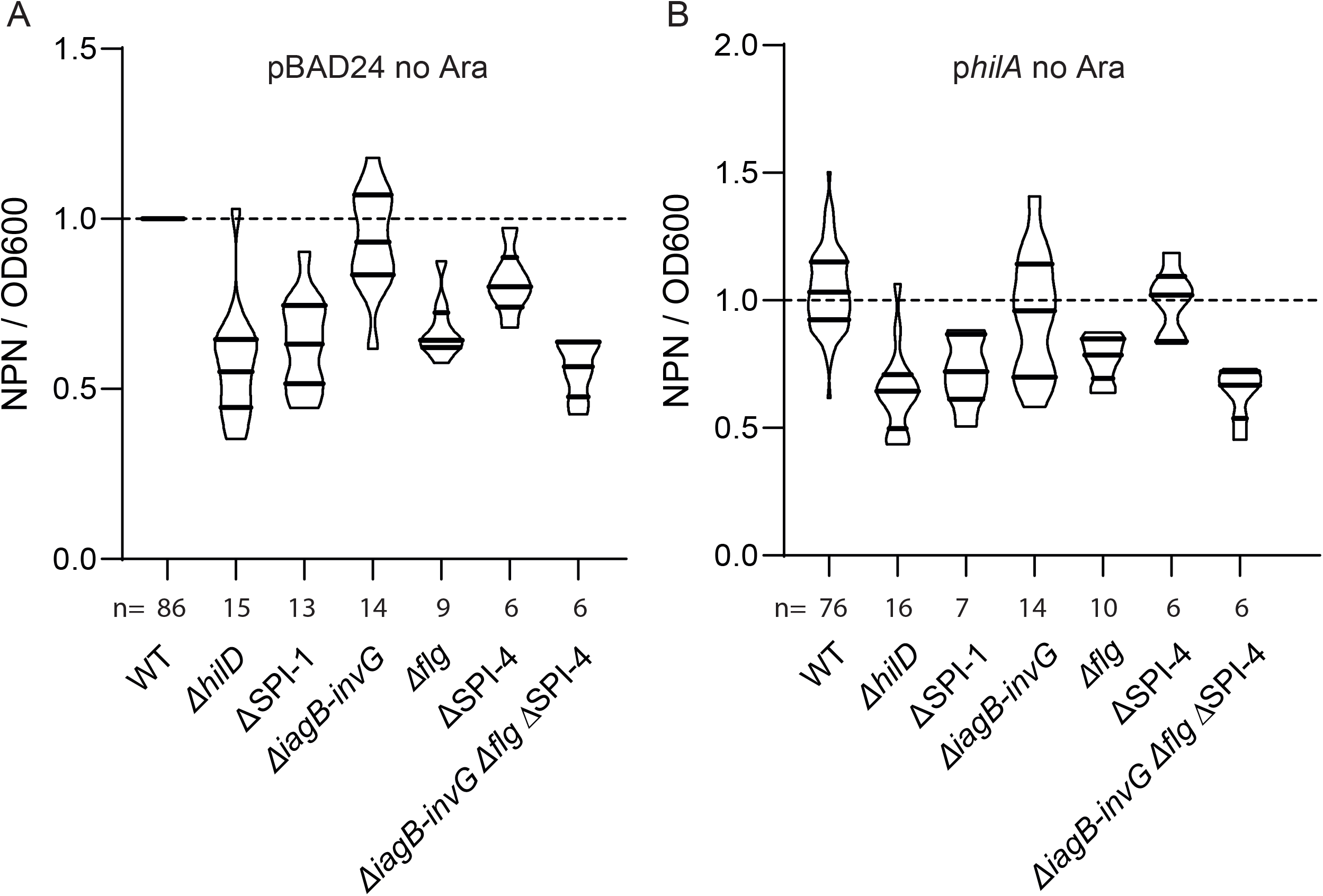
Experiments controlling NPN uptake in the absence of arabinose (corresponding to figure 1). Fluorescence of populations stained by NPN, corrected by the optical density at 600nm (OD600). **A**. Strains harboring pBAD24. **B**. Strains harboring p*hilA*. The **table S2** lists p values calculated for relevant comparisons. For comparisons against the WT, p values were calculated on the raw data using paired Wilcoxon tests. For comparisons between mutants, p values were calculated using the normalized data and unpaired Mann-Whitney tests.

**Fig. S3:**
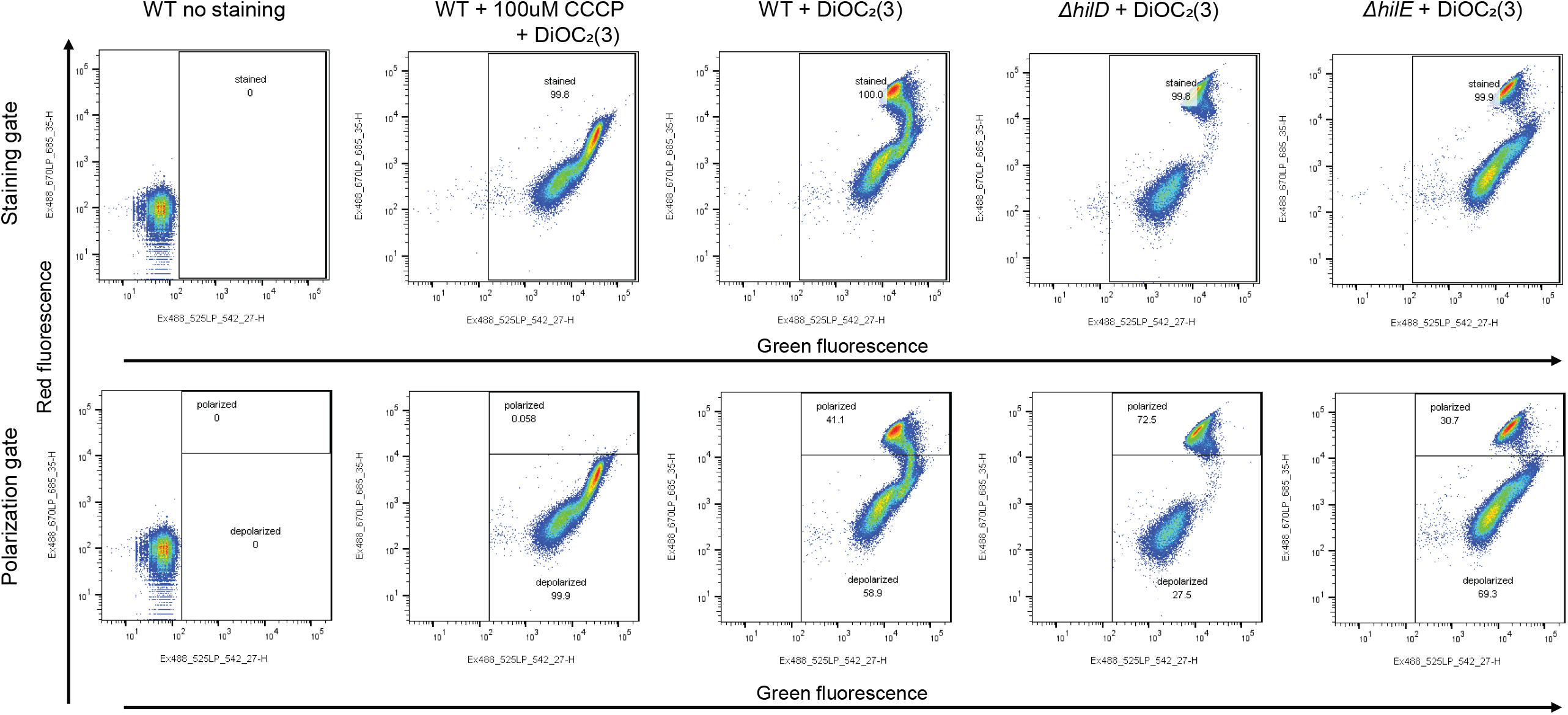
Gating strategy to determine the proportion of depolarized cells using DiOC^2^(3) staining. Flow cytometry plots showing events positioned according to intensities of their red (y-axis) and green fluorescence (x-axis) signals. Unstained WT control allowed delimiting the staining gate. Fully depolarized WT cells treated with 100 µm CCCP and 30 µm DiOC_2_(3) were used to delimit the depolarized cells among the stained cells. This gate has been used in all independent repetitions to calculate the fraction of depolarized cells in different *S*.Tm populations (**Fig. 2**).

**Fig. S4:**
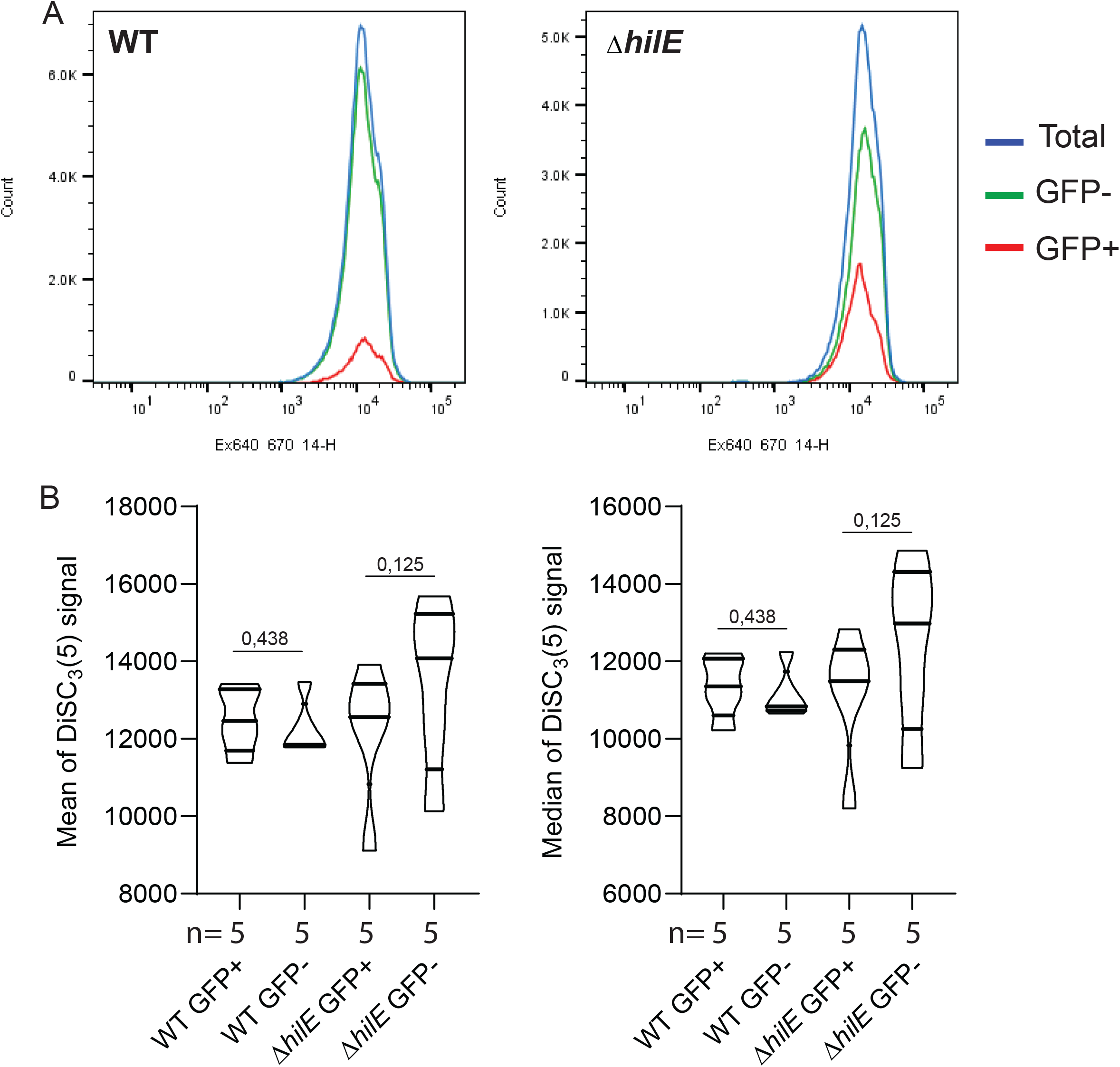
Membrane polarization in the absence of stress revealed with DiSC_3_(5). **A**. Red fluorescent signal in HilD expressing cells (GFP+) from WT (right histogram) or Δ*hilE* (left histogram) carrying a reporter P*prgH::gfp* (representative experiment). **B**. Mean (left) and median (right) of the red fluorescent signal due to accumulation of DiSC_3_(5) in WT or Δ*hilE* total, GFP+ or GFP-populations. p values were calculated on the raw data using paired Wilcoxon tests.

**Fig. S5:**
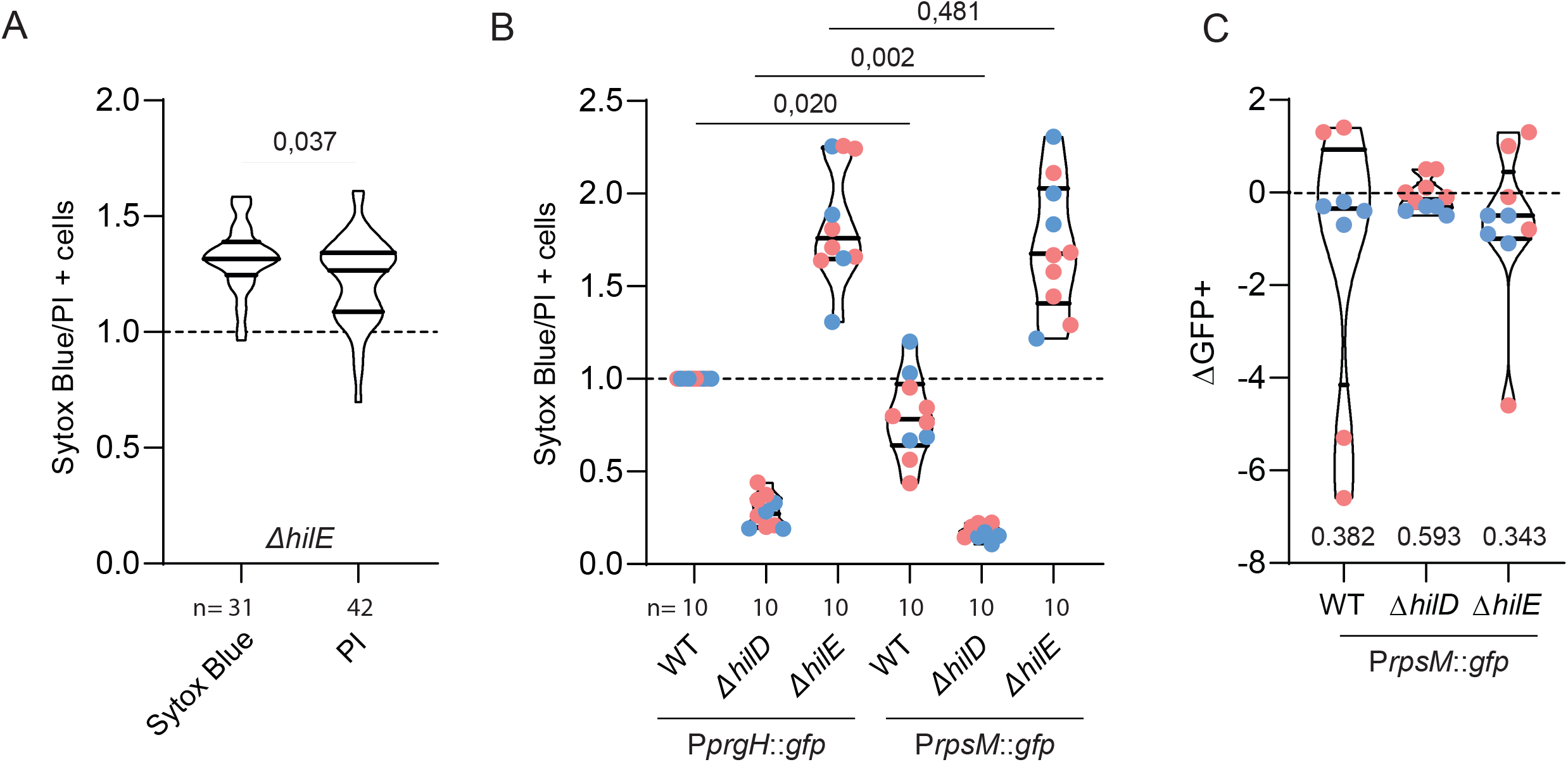
Validation of the cytometry analysis of cells exposed to a lethal concentration of Tris EDTA. **A**. Normalized frequency of Δ*hilE* reporter cells stained with either sytox blue or propidium iodide (PI) after treatment with 100mM Tris-10mM EDTA. **B**. Normalized frequency of cells stained with either sytox blue (blue dots) or PI (red dots) after treatment with 100mM Tris-10mM EDTA measured by flow cytometry. The graph shows the results for WT, Δ*hilD*, Δ*hilE* carrying a reporter P*prgH::gfp* or pM965 plasmid P*rpsM*::*gfp* (constitutive GFP expression). Values obtained with the WT reporter strain were used for normalization of the dataset. For comparisons against the WT, p values were calculated according to the raw data using paired Wilcoxon tests, comparisons between mutants or conditions p values were calculated using the normalized data and unpaired Mann-Whitney tests. **C**. Reduction of the GFP positive fraction (ΔGFP+) among WT, Δ*hilD* or Δ*hilE* alive cells (negative for sytox blue (blue dots) or PI (red dots)) treated with 100mM Tris-10mM EDTA compared to distilled water control. Significance of the deviation of the median from 0 estimated by Wilcoxon signed rank test.

**Fig. S6:**
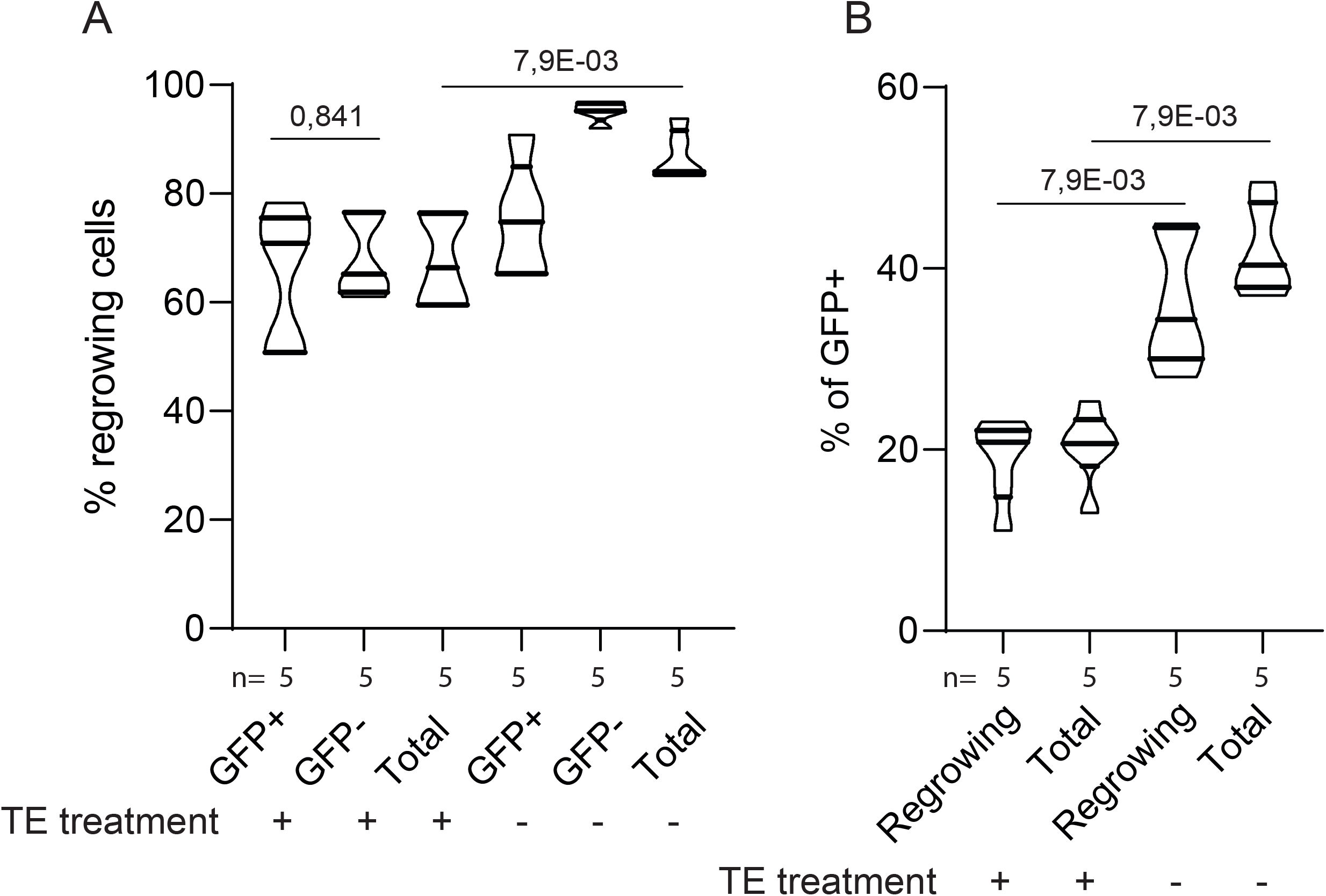
Time-lapse microscopy analysis of cells exposed to lethal concentration of Tris-EDTA and control. Time-lapse microscopy analysis of day culture of WT reporter strain (P*prgH::gfp*, GFP) treated with 100mM Tris 10mM EDTA and untreated control. **A**. Violin plots showing the fraction of cells able to form microcolonies among cells expressing the HilD regulon (GFP+) or not (GFP-) and in the total population. **B**. Fraction of GFP+ cells in regrowing and total population after treatment with 100mM Tris-10mM EDTA or untreated control. The p values were calculated using unpaired Mann-Whitney tests on raw data.

**Fig. S7:**
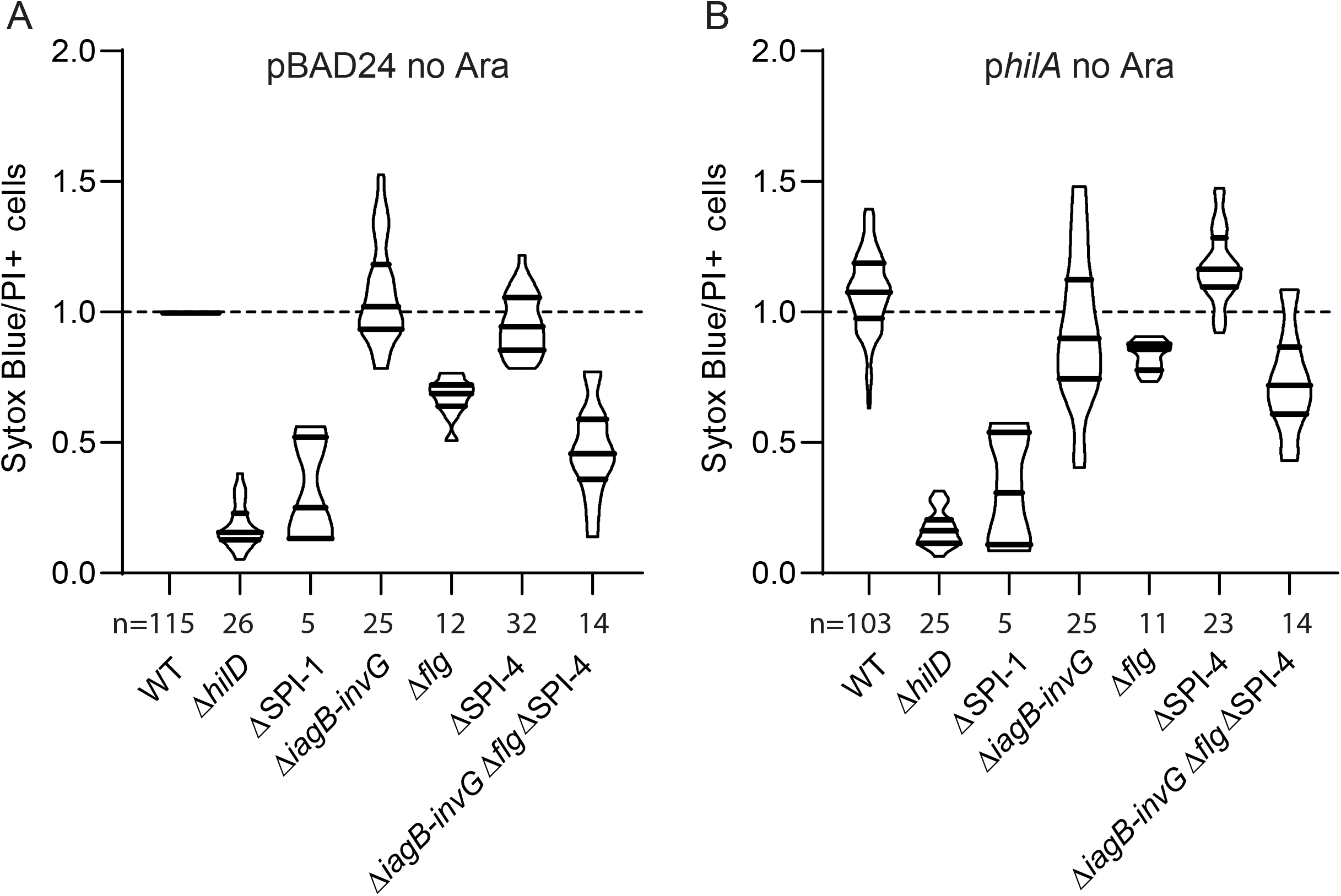
Experiments controlling death after exposure to lethal concentration of Tris/EDTA in the absence of arabinose (corresponding to figure 2). Cytometry analysis of cells stained by Sytox blue or PI after exposure to 100mM Tris-10mM EDTA. Data normalized by WT pBAD24. **A**. Strains harboring pBAD24. **B**. Strains harboring p*hilA*. The **table S4** lists p values calculated for relevant comparisons. For comparisons against the WT, p values were calculated on the raw data using paired Wilcoxon tests. For comparisons between mutants, p values were calculated using the normalized data and unpaired Mann-Whitney tests.

**Table S1 Proteomic analysis of the HilD regulon on sorted *S*.Tm cells**

**Table S2 Statistical analysis figure 1D, E and S2A, B**

**Table S3 CFUs**

**Table S4 Statistical analysis figure 2H, I and S7A, B Table S5 Strains and plasmids**

**Table S6 Primers**

